# Single cell DNA methylation ageing in mouse blood

**DOI:** 10.1101/2023.01.30.526343

**Authors:** Marc Jan Bonder, Stephen J. Clark, Felix Krueger, Siyuan Luo, João Agostinho de Sousa, Aida M. Hashtroud, Thomas M. Stubbs, Anne-Katrien Stark, Steffen Rulands, Oliver Stegle, Wolf Reik, Ferdinand von Meyenn

## Abstract

Ageing is the accumulation of changes and overall decline of the function of cells, organs and organisms over time. At the molecular and cellular level, the concept of biological age has been established and biomarkers of biological age have been identified, notably epigenetic DNA-methylation based clocks. With the emergence of single-cell DNA methylation profiling methods, the possibility to study biological age of individual cells has been proposed, and a first proof-of-concept study, based on limited single cell datasets mostly from early developmental origin, indicated the feasibility and relevance of this approach to better understand organismal changes and cellular ageing heterogeneity.

Here we generated a large single-cell DNA methylation and matched transcriptome dataset from mouse peripheral blood samples, spanning a broad range of ages (10-101 weeks of age). We observed that the number of genes expressed increased at older ages, but gene specific changes were small. We next developed a robust single cell DNA methylation age predictor (scEpiAge), which can accurately predict age in a broad range of publicly available datasets, including very sparse data and it also predicts age in single cells. Interestingly, the DNA methylation age distribution is wider than technically expected in 19% of single cells, suggesting that epigenetic age heterogeneity is present *in vivo* and may relate to functional differences between cells. In addition, we observe differences in epigenetic ageing between the major blood cell types. Our work provides a foundation for better single-cell and sparse data epigenetic age predictors and highlights the significance of cellular heterogeneity during ageing.

**Highlights:**

- Model to estimate DNA methylation age in single cells
- Large multi-omics dataset of single cells from murine blood
- Epigenetic age deviations from chronological age are greater than technical expected from technical variability
- Number of genes expressed increases with chronological and epigenetic age

## INTRODUCTION

The observation that specific DNA methylation patterns at CpG sites (hereafter DNAme) correlate robustly with age has led to the development of statistical models trained to predict age using DNAme measurements at a few hundred genomic loci (Bocklandt et al., 2011; Hannum et al., 2013; Horvath, 2013). These models, termed epigenetic or DNAme clocks, are capable of great accuracy, demonstrating their importance in the ageing field (Horvath, 2013). DNAme clocks have been created for a variety of species including human, mouse (Petkovich et al., 2017; Stubbs et al., 2017; Wang et al., 2017), and more recently across multiple species, demonstrating the evolutionary conservation of the DNAme ageing process (Lu et al., 2021). Noteworthy, while these model are trained based on chronological age, they do predict biological age, i.e. the biological functional decline associated with ageing, which is not always related to chronological age but rather also disease or disease risk (Belsky et al., 2020; Levine et al., 2018; Lu et al., 2019). These results illustrate the medical utility of DNAme ageing clocks and their relevance for measuring the effects of perturbations, such as drugs and other interventions. However, many DNAme - age associations are potentially confounded by changes in cell composition, particularly in blood where proportions of white blood cells are known to vary with age (Jaffe and Irizarry, 2014). Additionally, within a given tissue or cell type, it is not known whether DNAme ageing is a cell intrinsic process, such that all cells age at the same rate, or whether the ageing process is inherently heterogeneous and as such a cell population phenomenon. This underscores the need for better understanding and the relevance to develop single-cell age predictors and analysis. Additionally, the potential to use these age predictors in genetic or drug perturbation assays can facilitate massively parallel single-cell screens with biological age as a readout.

The possibility to profile DNAme in single cells genome-wide has become available in the last few years (Farlik et al., 2015; Gravina et al., 2016; Luo et al., 2017; Smallwood et al., 2014), but the datasets generated by these methods differ greatly from those generated by bulk approaches in three key aspects. Firstly, single-cell DNAme data is almost entirely binary, i.e. methylation values are either 0 or 100% for any given cytosine. Secondly, the datasets are very sparse with typically >90% missing values. Finally, the genomic coverage is essentially random, such that a dataset of a few hundred cells will contain no or only very few genomic loci with information from every single cell. These properties represent major challenges for generating age-predictors using current approaches which have relied on datasets with deep coverage at a consistent set of CpG sites.

The method we have developed here addresses these challenges by taking a different approach to traditional epigenetic clocks. Briefly, our method uses bulk DNAme datasets as a training set and generates a reference age profile for CpGs within our predefined age range using generalised linear models between age and DNAme level. The loci that showed the strongest age correlation in the bulk dataset and are present in a given cell are used to predict age in that cell. For a given locus, the observed DNAme state is compared to the expected states in the pre-calculated reference profiles and are transformed into probabilities that reflect the likelihood that an observed DNAme profile matches a specific age. The most likely age is then computed by multiplying the probabilities across loci and selecting the maximum. A recently published study used a similar approach (Trapp et al., 2021), and demonstrated the ability of their model to assign DNAme age to a small dataset of publicly available single cell adult mouse hepatocyte DNAme samples (Pearson r = 0.95, median absolute error = 2.1 months), but could not validate their model over a broader range of ages and cell types.

In the current study we overcome these limitations by generating a large single-cell dataset from peripheral blood cells of mice spanning a broad range of ages (10-101 weeks), using scM&T-seq (Angermueller et al., 2016) to profile single-cell methylomes and transcriptomes from the same cells. We show that our improved model allows us to more accurately predict epigenetic age in single cells and allow the model to be applied to (sparse) bulk methylation data. Our analyses also show that the epigenetic age variation of individual single cells at a given chronological age is wider than expected from technical noise and seems to be different across major blood cell types. Overall, we present a robust single-cell age estimator in the mouse and develop the methodological foundation for better epigenetic age predictors.

## RESULTS

### Ageing related changes in expression and DNA methylation in blood derived single cells

We collected peripheral blood from mice spanning ages from 10 to 101 weeks (Fig 1A, Suppl Table 1). Following red blood cell lysis, we FACS sorted individual cells into 96-well plates containing scM&T lysis buffer. The FACS gates were chosen to collect live and nucleated single cells, but otherwise not biassed for size nor granularity of the cells. Subsequently we performed scM&T-seq and generated paired single-cell methylomes and transcriptomes from 1,055 cells of which 853 passed methylation quality control (QC) filters, 981 passed expression QC filters and 823 passed filters on both (Suppl Table 1).

**Figure 1:**
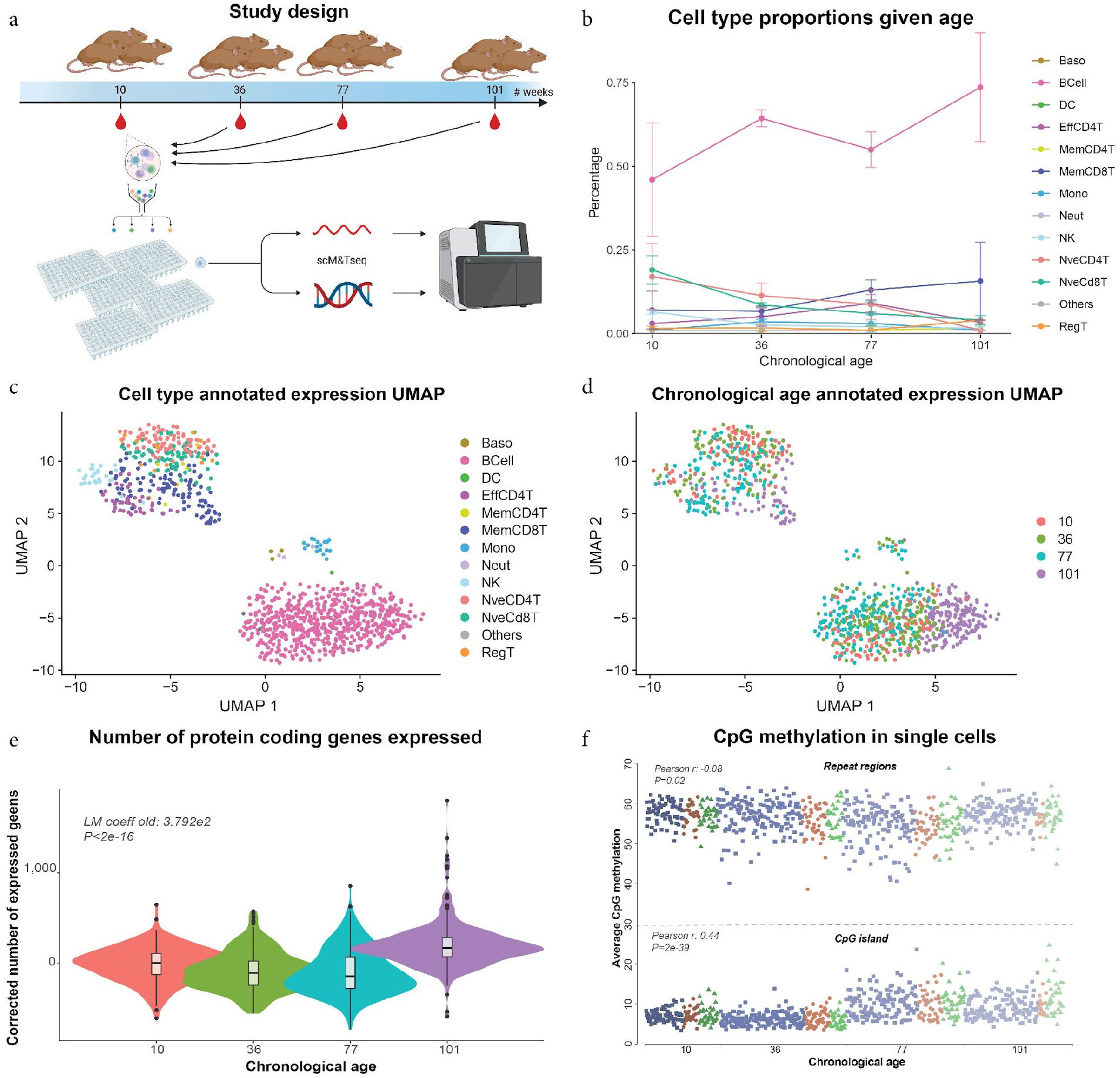
Study overview and data exploration. **a**) Illustration of the data collected for this study. We collected blood at 4 time points (10, 36, 77 and 101 weeks of age) from three mice each, FACS sorted single cells into 96-well plates and performed sc-M&T seq on 1,055 cells in total. **b**) Cell type composition overview stratified by chronological age. Lines depict the average percentages, and the error bars represent the variation (SD). **c-d**) Exploratory UMAP figures of the single cell expression data: **c**) Cell type annotated UMAP, and **d**) UMAP annotated by chronological age. **e**) Violin plots showing the number of genes expressed/detected in each cell at the different mouse ages. **f**) Average single cell DNAme levels in repeat regions (top half) and CGIs (bottom half). Cells are coloured by cell type (blue B-cell, orange CD4+ T-cell, green CD8+ T-cell) and ordered by age.

As a first step we performed a cell type annotation based on the single-cell transcriptome data. We leveraged a combined de novo and reference-based cell type annotation strategy (see methods for details). This resulted in a cell type classification and identification of the major categories of blood cell types. The cell type proportions reflect the expected distribution of B-, T, and other blood cells (Fig 1B) (Teo et al., 2021). Examining the cell type proportions as a function of age, using propeller (Phipson et al., 2022), did not show any significant ageing effect. Suggestively, we observe an increase in the B-cells proportion with age, agreeing with prior reports (Teo et al., 2021), but this change was not significant in our sample set, probably due to the lower sample and cell numbers.

Dimensionality reduction of the single cell transcriptome data showed clear segregation by cell type and not by age or animal (Fig 1 C&D). The DNAme data did not show clear separations by either age, cell type, or animal (Suppl Fig 1). This is likely confounded by sparse CpG coverage of the single cell data (Fig 1E). While we obtained data for approx. 1 million CpG sites in each cell, the number of CpGs covered between cells did drop markedly when jointly analysing multiple cells, even when we analysed the data at feature level (for instance at promoters or enhancers). The global analysis on gene expression highlights that there is a significant difference between the number of genes expressed in samples from younger ages (10, 36, and 77 weeks) verses from older mice (101 weeks; p<2e-16; Fig 1E), driving the separation observed in the UMAP plots (Fig 1C&D). This effect is consistent in the three major cell types (Suppl Fig 2A-C), and we were able to replicate the effect in the three tissues of the Tabula Muris Senis study (The Tabula Muris Consortium et al., 2020; Suppl Table 2). Additionally, we replicated the effect in a large human PBMC cohort, the OneK1K dataset (Yazar et al., 2022), where we also find that samples below 55 years of age (roughly equivalent to under 77 weeks in mouse), have significantly fewer genes expressed (p<2e-16; Suppl Fig 2D) as compared to samples over 64 (roughly equivalent to over 101 weeks of age in mice).

When comparing cells at the level of DNAme at repeat regions or CpG islands (CGIs; Fig 1F) we observe limited variation between the cells over cell types and age indicating high quality data (see methods for details). Interestingly, at shared CGIs, DNAme levels increased with age (r=0.44, p<2e-39), a phenomenon that could be replicated using data from sorted blood cells (r=0.86, p<9e-10, see methods for details, Suppl Fig 3A). In contrast, in repeat regions we observed a weak but significant negative correlation with age in our single cell data (pearson r=- 0.08, p=0.02, see methods for details), and this effect was even stronger in the bulk data sets (pearson r=-0.45, p=0.01, see methods for details, Suppl Fig 3B).

We next assessed transcriptional differences at individual genes, using MAST (see methods for details), and DNAme differences at the level of individual enhancers and promoters using a generalised linear mixed model (see methods for details). Interestingly, at per-gene level we did not see any significant transcriptional changes in CD4+ T cells, and only three genes that were significantly differentially expressed in CD8+ T cells (FDR < 10%, Suppl Table 3A). Within B cells we identified 97 genes with significant changes with age (FDR < 10%, Suppl Table 3B). Given that we observed an increase of genes expressed after 77 weeks of age we also tested for differences between cells from the three youngest ages and the oldest age. We found one gene in CD4+ T cells (Suppl Table 3C), and two genes, which were also found in the quantitative analysis, in CD8+ T cells (Suppl Table 3D). Within the B cells we found additional 57 genes that were exclusively significantly differentially expressed between the three youngest ages and 101 weeks (Suppl Table 3E). The majority of these genes (30) were in the positive direction in the discrete part of the MAST model. Matching our prior observation, that a larger number of genes are expressed in older ages. 66 of the 97 global age associated B-cell genes increased in expression level with age, and among this set we found a strong enrichment for ribosomal RNA processes (Suppl Table 3D & 4). This is in line with previous studies in blood transcriptomic changes (Frenk and Houseley, 2018; Teo et al., 2021). We next assessed age-related DNAme changes in both enhancers and promoters, and identified 3 enhancers, all in CD4+ T cells, and 48 promoter regions, 22 in CD4+ T cells and 27 in CD8+ T cells, that had age associated changes (Suppl Table 5).

### Modelling epigenetic ageing in single cells

#### Training the model on bulk data

We next set-out to quantify the difference in epigenetic ages within and between the different timepoints. To do so we built a DNAme ageing clock (scEpiAge) to predict ages of individual cells. In contrast to classical epigenetic ageing models, we leveraged an approach that is flexible enough to overcome differences in coverage and can deal with a high degree of sparsity, thereby making it applicable to single cell as well as sparse bulk data. Specifically, we built two models, one for blood (scEpiAge-blood) and one for liver (scEpiAge-liver). We combined newly generated bulk DNAme data (n=226), which included sorted blood cell types (n=32) and liver samples (n=33), [new Babraham dataset; see methods] with five publicly available bulk data sets (Meer et al., 2018; Petkovich et al., 2017; Reizel et al., 2015; Stubbs et al., 2017; Thompson et al., 2018), totalling 411 samples (Suppl Table 6, 262 in blood and 138 in liver) and age ranges from 1 to 153 weeks (3 to 153 weeks for blood, and 1 to 128 weeks of age for liver).

Our scEpiAge setup, illustrated in Fig 2 A&B, is similar to the genetic and epigenetic distance based prediction methods used in other settings (Eirola et al., 2013; Han et al., 2020a, 2020b; Karimzadeh and Olafsson, 2019) and also by Trapp et al (Trapp et al., 2021), but includes the following major modifications: (1) we generalised it to work on both bulk and single-cell data, (2) we used a significantly larger number of reference samples (2.3 times more samples for liver and 3.6 times more samples for blood) originating from five different studies, and (3) we optimised feature selection and modelling. In short, we selected per tissue the top age correlating sites pruned to only capture independent age associated CpGs. The sites that were pruned away are held in reserve to be used as backup sites in case the CpG with the strongest age signal is not measured. Next, we fitted a binomial regression model per CpG site modelling the relation between methylation and age. Using this fit we calculated the expected methylation level of a CpG at a given age. To predict the age in a new bulk or single cell sample we summed the difference between the observed methylation state of the sites covered in the new sample, and the expected methylation levels at each of the ages. The age of the new sample is given by the expected DNAme profile that is closest to the new methylation profile (Fig 2B). More detailed information can be found in the methods.

**Figure 2:**
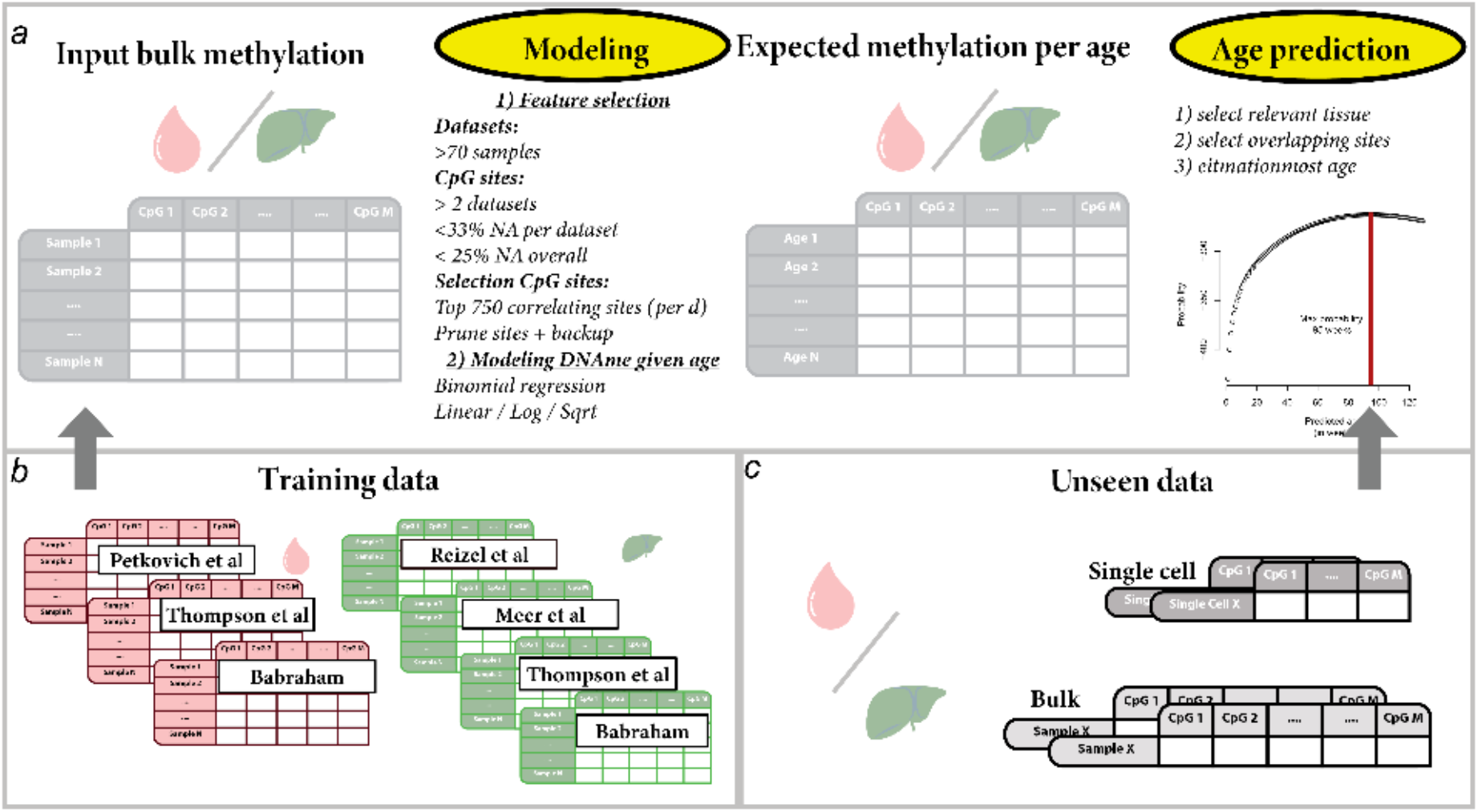
Overview of the sc-EpiAge modelling setup. **a-c**) Schematic representation of the scEpiAge model during training, testing and application. **a**) Modelling setup going from input data (left), via feature selection and age modelling (left centre), to an expected DNAme given age matrix (right centre), and to application for age prediction (right). **b**) Datasets used to train the two scEpiAge models: We selected DNAme datasets from blood and liver, which we generated in this study or obtained from from five published studies (Stubbs et al., 2017); (Petkovich et al., 2017); (Reizel et al., 2015); (Meer et al., 2018); (Thompson et al., 2018). **c**) Depiction of unseen data either in the validation stage or after model building.

To find the optimal models we leveraged ten-fold cross validation (methods). Here we tested the transformation of sites (linear, log, square root), the selection of the most age correlating sites and the number of sites (ranging from 50 to 2,000). Our best performing model leveraged information from 750 sites, selected the best transformation function based on model fit per CpG site, and selected the top associated sites based on the average relation to age over the datasets. The best scEpiAge model has a cross validation median absolute error (MAE) of 8.2 weeks of age in blood and a MAE 3.9 weeks in liver. We compared our modelling setup to a direct reimplementation of the Trapp et al. model (Trapp et al., 2021), where the respective MAE is 16 weeks in blood, and a MAE of 7 weeks in liver (Methods and Suppl Fig 4). When applying the model to the whole bulk training dataset we obtain a final MAE of 8.2 in blood and 4.6 in liver (Fig 3 A&B). Using left out validation samples, we could validate these errors, which we found to be in line with the error on unobserved data (Fig 3 C&D; MAE: 5 for liver and MAE: 10 for blood).

**Figure 3:**
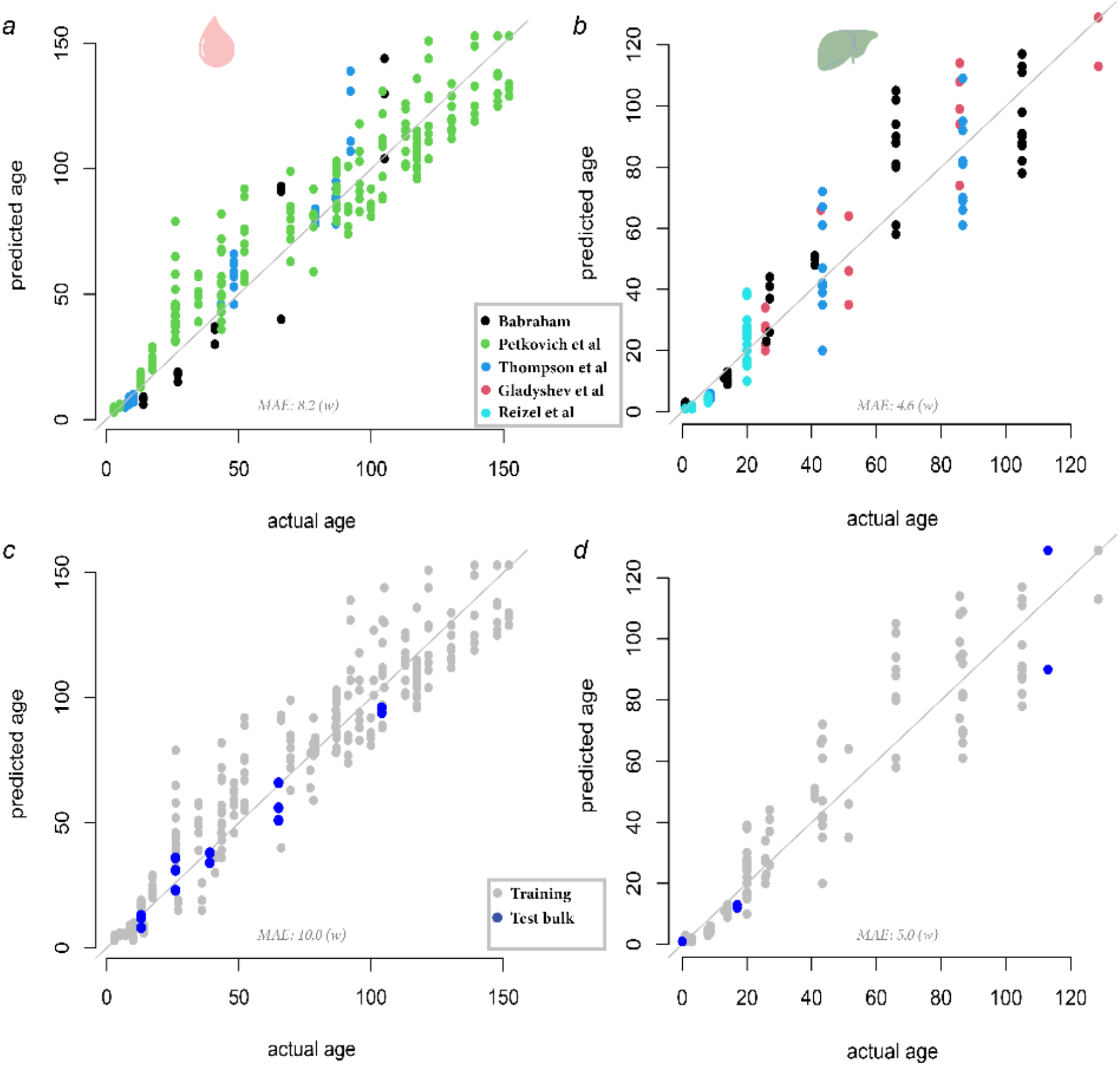
DNAme based age prediction performance in bulk. **a-d**) Performance of the model on training data and on simulated single cells. Performance of the models on the training set in blood **a**) and liver **b**); performance of the models on independent bulk blood **c**) and liver **d**) data.

To gain insight into what types of sites are important for the scEpiAge models we performed site enrichments, including the back-up sites, as well as for the Stubbs et al. multi-tissue clock sites (Stubbs et al., 2017) (Suppl Fig 5). We find that the enrichments for the main scEpiAge models, without backup sites, and the models including backup sites have the same significant enrichment regions, and only marginally different enrichment statistics. There is only one class of features, i.e. coding sequences in the scEpiAge-blood model, that is not significantly enriched when not taking into account the backup sites, but the enrichment statistic is very similar in both settings. This shows that the putative biologically relevant effects captured by the main and backup sites are highly complementary. In general, we find that the scEpiAge-liver model selects similar regions as compared to the Stubbs et al. multi-tissue model while the scEpiAge-blood model selects markedly different regions. In the scEpiAge-blood model we do not see the significant depletions for promoters, CGIs and 5’ UTR regions that we do observe in both the scEpiAge-liver and the Stubbs et al. multi-tissue model (Stubbs et al., 2017). It is worth noting that the Stubbs et al. multi-tissue model is built based on heart, lung, brain, and liver DNAme datasets, but does not include blood DNAme data. This could explain the discrepancy in site enrichments between the various modes.

Next, we compared the scEpiAge model to a classical epigenetic ageing model based on regression models. We decided to use the Stubbs et al. clock (Stubbs et al., 2017) as an example of this type of model and compared it to the new scEpiAge-liver model. For the comparison we only considered samples in the range of 1 to 41 weeks of age, to match the published model. We found that the original Stubbs et al. model (including normalisation) only works on samples from the original cohort (Stubbs et al., 2017) and the Reizel *et al*. study (Reizel et al., 2015), both samples that comprise the training and testing data of the Stubbs *et al*. clock. All other samples, i.e. Meer et al. (Meer et al., 2018) or Thompson et al. (Thompson et al., 2018), do not generate a prediction. However importantly the scEpiAge model can predict ages for samples from all studies. We then focused on the samples (n=63) that were part of the training sets in both models. The MAE for these samples in the Stubbs et al. model is 0.8 weeks and 3 weeks in the scEpiAge-liver model. The correlation between the predictions of the two models is high (pearson r^2^=0.87), but lower than each one individually to chronological age (pearson r^2^: 0.92 (Stubbs et al., 2017), and pearson r^2^=0.88 (scEpiAge-liver)).

#### Assessing the performance of the model on simulated single cells

Next, we assessed the performance of the model on single cells. We first simulated 50 single cell methylomes by downsampling bulk samples in the validation dataset (see methods). We found that the error on the simulated liver cells was 4.6 weeks of age, and in blood the error on the simulated single cells was 5.2 weeks of age (Fig 4 A&B). We compared the scEpiAge predictions with the predictions using the Trapp et al model, where we found that for the simulated single cells the liver model results in an MAE of 8.3 weeks of age and for blood an MAE of 10.0 weeks of age, an error increase of 3.1 weeks of age for liver and 1.7 weeks of age in blood (Suppl Fig 6A&B).

**Figure 4:**
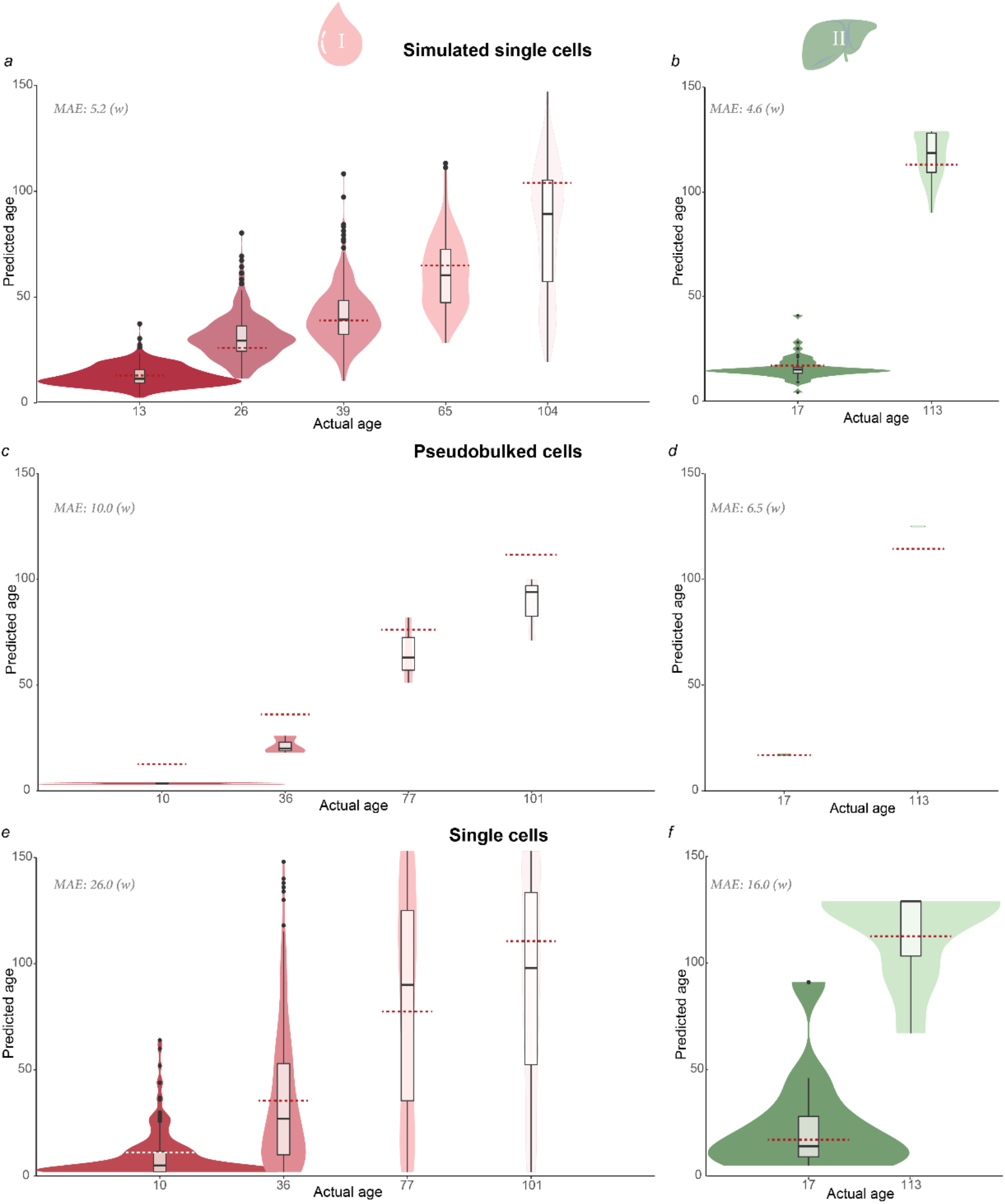
Predicting methylation age in simulated and real single cell data. The predictions for epigenetic age shown in this figure have been made using the scEpiAge model for blood or liver described in Figure 2. **a-b**) Predicted DNAme age of simulated **a**) blood or **b**) liver single cells. **c-d**) Predicted DNAme age of pseudo bulked real **c)** blood and **d**) liver single cell data. **e-f**) Predicted DNAme of the single cell **e**) blood and **f**) liver data.

#### Implementation and validation on single cells

Next, we moved to predicting age in real single cells, using our blood dataset together with published single-cell methylation data from hepatocytes (Gravina et al., 2016). First, we assessed the performance at pseudo bulk level, aggregating the cells by donor, to estimate the impact of batch effects and different sequencing methods on the epigenetic clocks. We found in blood a MAE of 10 weeks and in liver a MAE of 6.5 weeks, both in line with the errors in bulk but higher (Fig 4 C&D, Trapp et al. results are in Suppl Fig 6 C&D).

Observing close to expected age predictions in the pseudo bulk setting we moved to single cell predictions, where we found that the error of the scEpiAge models increased substantially. We can predict ages in all cells which pass QC, both in blood and liver, but the MAE increases to 26 weeks in blood and 16 weeks in liver when predicting real single cells (Fig 4 E&F), and correlation between expected age and real age is reduced (pearson r =0.6; p<2.2e-16). We compared our performance to the Trapp et al. model, where we observed similar prediction errors: 24.26 in blood, 14.2 in liver (prediction rates from the Trapp et al. model 98% of the blood cells and 100% of the liver cells) (Suppl Fig 6 E&F). We observed a relationship between the age prediction and the coverage of the cells in blood (r=-0.36, p<2.2e-16) indicating that the number of covered sites influences our ability to accurately predict age. Based on a simulation we noted that we need at least 40 covered sites, at which point in the simulated cells the mean error is below 1 week of age (see methods, Suppl Fig 7). When focusing on cells fulfilling this requirement, we retain 72.5% of all cells in our single cell blood data (95% of cells in the liver data), and the MAE decreases to 20 weeks in blood and 15.5 weeks in liver. For the Trapp et al. model we observed a similar but smaller gain (MAE in blood 21.8 weeks). The spearman correlation between our model and the Trapp et al. prediction in blood is 0.51 (spearman rho, Suppl Fig 8a), while in liver the correlation is 0.75 (spearman rho, Suppl Fig 8b).

### Epigenetic age predictions in blood at single cell resolution

Next, we focused on the data we generated from blood cells spanning a wide range of ages and predicted DNAme age for each cell (Fig 4E). When taking a closer look at these predictions we observed that the median scEpiAge predictions of cells at a given age was very similar to the expected age, i.e. at 10 weeks of age the mean prediction was 8.4, at 36 weeks of age it was 36.2, at 77 weeks of age it was 83 and at 101 weeks of age it was 88. The variance increases with age, with a standard deviation of 10.2 at 10 weeks, 32.7 at 36 weeks, 45 at 77 weeks, and 42 at 101 weeks. We asked if the observed epigenetic age predictions and the individual cellular deviation from the organismal (chronological) age reflect potential biological differences or result from possible technical noise of the model. We leveraged cell specific simulation, based on the expected age and CpG sites covered in the cell of interest, to find outliers (see methods). We found a total of 120 cells (19.4%) with a significant deviation from the expected age (Fig 5B; empirical FDR 5%), i.e. cells where the predicted epigenetic age deviated significantly more from the expected (chronological) age than technically expected. Most of the significant deviations were cells that were significantly younger than expected (84 out of 120). This is in line with the error that shows more under predictions as compared to over predictions (Fig 5A-B). Most of the extreme predictions were found at 36 weeks of age, where 24% of the predictions were significantly deviating from the expected age, of which 20% were predicted to be younger and 4% predicted to be older.

**Figure 5:**
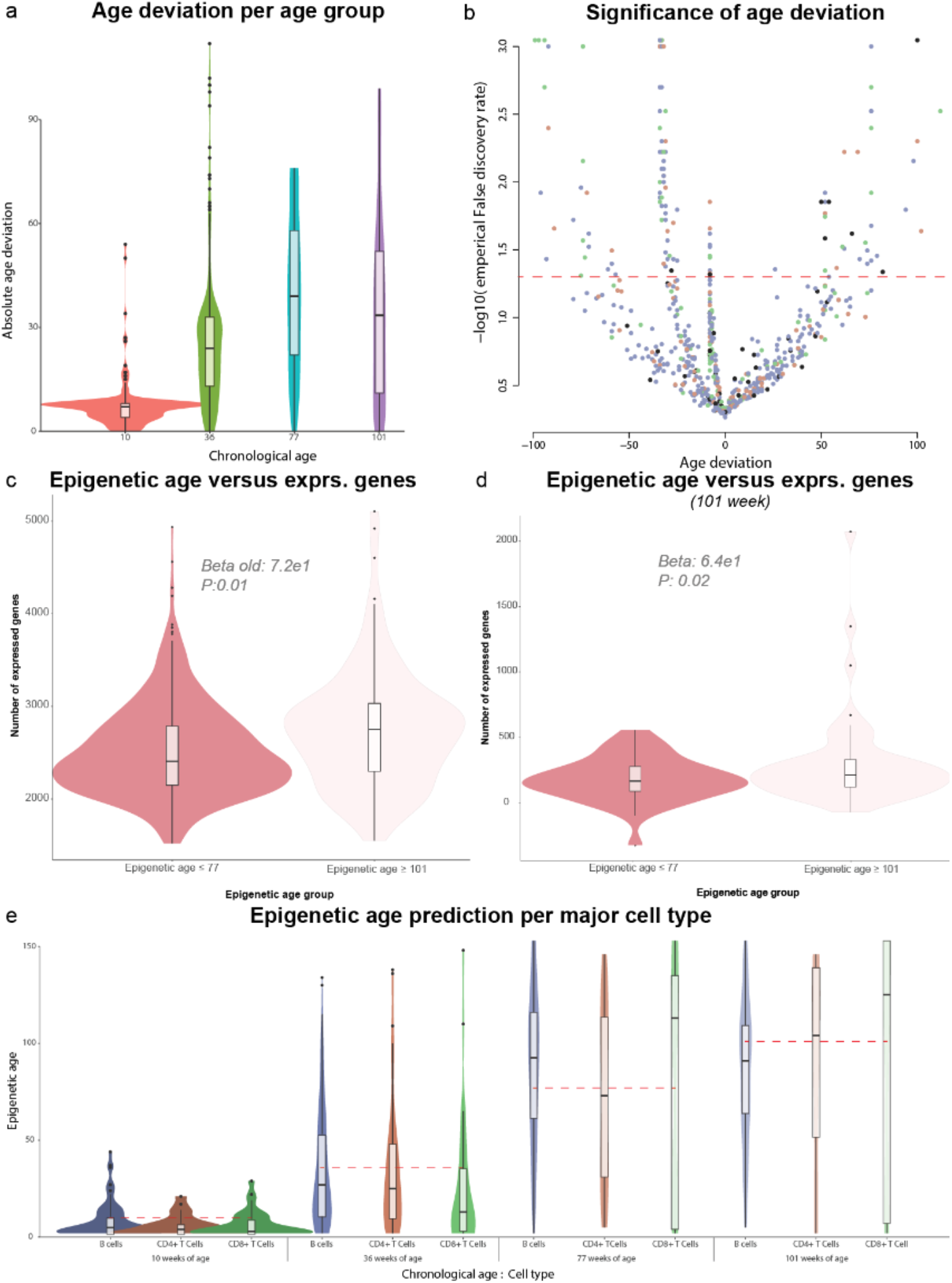
Epigenetic age predictions in blood at single cell resolution. **a**) We calculated the deviation between the real chronological age and the DNAme age predictions of our scEpiAge model for each single cell. Depicted are the deviations for each age group. **b**) For each cell we estimated the empirical false discovery rate and plotted these against the age deviations between chronological and DNAme age. Individual cells are coloured by cell type: blue B-cell, orange CD4+ T-cell, green CD8+ T-cell, black other. **c**) Epigenetic age predictions per major cell type, grouped by chronological age. **de**) Number of expressed genes in cells with an DNAme age below 77 weeks or DNAme age above 101 weeks, showing cells from **d)** all chronological age groups or **e)** only from chronological age 101 weeks.

We then asked if the scEpiAge differences are associated with global and specific gene expression effects. As observed for chronological age we observed a significant increase (LM p=0.1) in the number of expressed genes in older cells (epigenetic age ≥ 101 weeks) versus younger cells (epigenetic age ≤ 77 weeks) (Fig 5C). Given the relation between epigenetic age and chronological age we also tested this within each chronological age, and corrected the number of expressed genes for the number of sequenced reads, where we could also significantly identify this effect in the 101 week old cells (LM p=0.02) (Fig 5D). In other chronological age groups we could not replicate the effect, however the number of old cells (epigenetic age ≥ 101 weeks) in these categories is low. When replicating effects from chronological age we observe that we can replicate three of the 97 ageing associated genes in B-cells (Suppl Table 2B). In a subsequent genome-wide analysis we did not find any significant changes in B-cells or CD4+ T-cells. In the CD8+ T-cells we identified 3 genes that were differentially expressed with ageing (FDR <10%), i.e. *Ttn, Rpl36a* and *Txnip*.

Interestingly, when splitting the scEpiAge predictions per major cell types, i.e. B-cells, CD4+ and CD8+ T-cells, and others, we observed that the predictions for CD4+ and CD8+ T-cells were in general lower as compared to B-cells. On average, CD8+ T-cells are estimated to be 3.5 weeks younger than B-cells, CD4+ T-cells are found to be 0.5 weeks younger than B-cells. Of note, the model is overall underestimating epigenetic age by 1.3 weeks compared to chronological age) (Fig 5E). We also observed that a large number of T-cells had significantly younger age predictions (25.3% of CD8+ T-cells and 15.6% of CD4+ T-cells, versus 11.6% of B-cells), but T-cells also had more extreme age predictions on the other hand, with 8.4% of the CD8+ T-cells estimated to be significantly older than expected (5.8% of the CD4+ T-cells), versus only 4% of the B-cells(Fig 5B-E). To verify these findings, we investigated age predictions of sorted blood cells, i.e. CD4+ T-cells, CD8+ T-cells, and B-cells, and observed that in bulk CD4+ and CD8+ T-cell data ages were consistently lower as compared to the B-cells from the same mouse which is in line with our single cell data.

## DISCUSSION

Ageing at the organismal or tissue level is characterised by progressive functional decline and deterioration, increasing the risk of death (López-Otín et al., 2023, 2013). Unicellular organisms show signs of ageing and are generally not immortal (Florea, 2017), however our knowledge of how individual cells age within multicellular organisms remains poorly understood. Several scenarios are plausible to explain the age-related changes of tissues, of which the extremes would be that 1) all cells age at a uniform rate and continuously lose their functional capacity, or 2) cells could only have 2 possible age-states, i.e. young and old, and the age of a tissue is determined by the relative proportion of young and old cells. Ageing at the single cell level could also be a combination of these two extreme scenarios, where cells age at different rates and functional decline occurs at varying rates. Addressing these fundamental questions has remained challenging and despite several studies assessing transcription in single cells at varying ages (The Tabula Muris Consortium et al., 2020; Zou et al., 2021), we cannot yet resolve this issue. The use of epigenetic, in particular DNA methylation based, age predictions has thus far shown to be the most accurate way to assess biological age (Hannum et al., 2013; Horvath, 2013; Stubbs et al., 2017) and this motivated us to develop two new scDNAme age predictors (scEpiAge-Blood & scEpiAge-Liver) to explore this concept in single-cells and generate a large single-cell multi-omics dataset, assessing DNA methylation and transcription from nucleated blood cells, from mice spanning a broad range of ages.

The development of a new single-cell age predictor was necessary, as single-cell data differs in key aspects from bulk sequencing data, being very sparse and primarily binary. In an earlier work, where muscle stem cells from young and old mice were studied (Hernando-Herraez et al., 2019), we addressed this issue by aggregating the single cell datasets and then retraining the original model (Stubbs et al., 2017), only including sites covered also in the single cell dataset. While this approach was feasible, it is not scalable and only applicable if several preconditions are fulfilled. Our new epigenetic ageing models (for liver and blood) address these challenges by building on bulk methylation datasets (newly generated ones and publicly available ones), from corresponding tissues, to generate a theoretical methylation profile for each age (and tissue type). Distinct from previous generations of epigenetic clocks (Horvath, 2013; Levine et al., 2018; Meer et al., 2018; Stubbs et al., 2017; Thompson et al., 2018), the scEpiAge model can be applied even when not all sites are present, lowering the performance but enhancing usability substantially. To further improve performance and usability we included the use of backup-sites, CpGs with a DNAme profile similar to and in close proximity to the main site. Additionally we show that our generalisations in the modelling of the expected DNAme profiles increases accuracy over multiple datasets and in complex tissues such as blood, relative to a recent similar clock model by Trapp et al. (Trapp et al., 2021). By leveraging *in silico* simulated single cell DNAme profiles matching the sites covered in a single cell and the DNAme profile at a given age, we can now add a confidence limit on how certain we are that a predicted age is out of the expected range. Lastly, our model can be applied to (sparse) bulk bisulfite data, for low depth long-read sequencing methods or also for large scale screening approaches, enhancing the usability of epigenetic age models on sequencing data in general.

When initially exploring our mouse blood single-cell multi-omics dataset spanning a broad range of ages, we found only minor age-related differences. Cell type proportions only showed a small but non-significant overall increase of B-cells with age (Fig 1B), in line with prior reports (Teo et al., 2021), and similarly we only observed minor transcriptional changes of the single cells at specific cell type level (Fig 1D). Nonetheless, several of these differentially regulated genes had previously already been associated with ageing (Teo et al., 2021), suggesting that some of the transcriptional changes do reflect age-related effects. Interestingly, we find a strong effect on the number of genes expressed with age, which we replicated in external datasets. However, the pattern is complex. The number of genes expressed first decreases minimally up to 77 weeks of age in our data, followed by a strong increase at our 101 week old time point. We designed our replication analyses to match this phenomenon and find that it replicates in several Tabula muris datasets and in a large-scale human PBMC dataset. However, to better dissect this phenomenon a denser sampling procedure would be required as well as more replicates.

Mirroring the results of the gene-expression analysis, we found only minor differences in the DNAme landscape with age (Suppl Fig 1). Interestingly, we found that DNAme levels at CGIs increased with age in the single cell dataset and in repeat regions we observed a weak but significant negative correlation with age (Fig 1F), and both results could be confirmed in bulk data from sorted blood cells. This result indicates that cell type specific global changes might be accumulated over time but have likely been masked due to cellular heterogeneity in past studies. These DNAme changes could be the result of imperfect DNAme maintenance, resulting in gain of DNAme at hypomethylated (CGIs) or loss of DNAme at hypermethylated (repeat) regions.

It is worth noting that, due to technological limitations of single-cell methylation methods, the number of cells we profiled is relatively large compared with published DNAme datasets yet small compared with single-cell RNA-seq datasets. As such, our transcriptional analyses are relatively low powered, and we may not be able to detect smaller age-related transcriptional differences in our study, and in particular small transcriptional effects caused by small DNAme changes in CGI promoters or repeats. Additionally, it is worth noting that DNAme datasets are inherently sparse which negatively impacts the power of our analyses.

Comparing the predicted ages with the known chronological ages we found that both pseudo-bulk datasets as well as the average age of all cells of a given age group agreed with the expected (chronological) age, showing the consistency of our new approach and prior models (Stubbs et al., 2017; Trapp et al., 2021). A comparison of our scEpiAge-liver and scEpiAge-blood predictor showed that epigenetic ageing signals are stronger in the liver as compared to blood, possibly reflecting the more complex cell type distributions in blood. It should also be noted that while we had more training data for the scEpiAge-blood model (than for scEpiAge-liver), these were primarily heterogenous and not cell type specific (FACS sorted) datasets, which made identifying a global, rather than cell type proportion affected age signal, more difficult and potentially leading to a less robust model. Alternatively, the differences could be explained by dataset and age span differences, where the scEpiAge-blood model works across a larger window and the data is less uniformly distributed over data sources and ages, which might negatively impact the modelling. We assume that these reasons combined explain that even with fewer samples in the liver (138 liver versus 262 blood), we were able to predict ageing in the liver more precisely as compared to blood (MAE 3.9 in liver versus 8.2 in blood).

A closer analysis of the single-cell scEpiAge predictions uncovered ageing heterogeneity which was larger than technically expected for 19.4% of the cells. This indicates that, according to scEpiAge, younger and older cells co-exist and strikingly even at older ages a pool of epigenetically young cells remains, in line with a recent in vitro study, which shows that the epigenetic age heterogeneity observed between cell clones from the same donor is greater than the variability of the clock prediction (Kabacik et al., 2022). In terms of transcriptional differences with age, we found differential expression of *Rpl36a*, a ribosomal gene which is down-regulated with higher scEpiAge, which is in line with our results on chronological age and the literature (Frenk and Houseley, 2018). Also *Txnip*, a major regulator of cellular redox signalling which protects cells from oxidative stress, was found to be significantly lower expressed at higher scEpiAges, and the loss of *Txnip* was shown to induce premature ageing in hematopoietic stem cells (Jung et al., 2016). Follow-up work will increase the sample number and hopefully thereby increase the statistical power to detect other relevant putative functional differences.

The fact that we also found most of the extreme predictions at 36 weeks of age, where 24% deviated from the expected age, (20% predicted to be younger and 4% predicted to be older), could suggest differences in ticking rates between cells at a given age, most pronounced in mature mice. Furthermore, our results showing that CD4+ and CD8+ T-cells were consistently younger when compared to B-cells from the same mouse, could indicate that T- and B-cell ageing differs, and ageing occurs on a different scale in T-cells compared to B-cells. Yet, these findings are based on a low number of donors and cells and need to be verified in a larger dataset.

In summary, we have developed a framework to expand our understanding of ageing, address key questions related to cell type specific age-related changes and also better understand the process of rejuvenation in single cells. The future applications are manifold and importantly, the new epigenetic age prediction concept can also be applied to other species, including humans.

## Methods

### Animals and Sample collection

Mice were bred and maintained in the Babraham Institute Biological Services Unit (BI BSU) under Specific Opportunistic Pathogen Free (SOPF) conditions. C57BL/6 J mice (supplied by Charles River Laboratories) were imported into the BI BSU by embryo transfer and bred there (/Babr) as a SOPF colony in plastic film isolators. After weaning, male mice were transferred to individually ventilated cages in groups of between two and five mice. All mice were fed CRM (P) VP diet (Special Diet Services) *ad libitum* and received sunflower seeds, poppy seeds or millet at cage-cleaning as part of their environmental enrichment. The health of mice was monitored closely and any mouse exhibiting clinical signs of ill-health or distress that persisted for more than three days was culled. In this way, all the mice maintained, remained ‘sub-threshold’ with respect to the UK Home Office severity categorisation. Any mice exhibiting any gross pathology upon post-mortem examination was excluded from this study.

Samples for bulk datasets were collected as described before (Stubbs et al.). All tissues were snap-frozen directly after isolation. Genomic DNA was isolated from ~10 mg frozen tissue using the DNeasy Blood & Tissue Kit (Qiagen).

Sorted blood cell types (CD4+, CD8+ T-cells, and B220+ B-cells) were collected from blood of the same animals used in the Stubbs et al study (Stubbs et al., 2017). Briefly, red blood cells were lysed, and the samples were then incubated with anti-mouse CD3 (clone 145-2C11), anti-mouse CD4 (clone GK1.5), anti-mouse CD8a (clone 53-6.7) and anti-mouse B220 (clone RA3-6B2). All antibodies were purchased from Biolegend, and used at a dilution of 1:300. Cells were then washed once in PBS with 0.5% BSA, and stained with DAPI. Cells were flow sorted (BD Aria III sorter) directly into RLT plus lysis buffer (Qiagen). Lymphocytes were gated based on forward and side scatter, then gated on singlets, followed by live cells. T-cells were selected by gating for CD3+ and then gating for CD4+ or CD8+. B-cells were selected by gating for CD3- cells and then gated for B220+ cells.

Blood samples for single cell analysis were collected and directly processed. Red blood cells were lysed using 2 rounds of red blood cell lysis (RBC Lysing Buffer, SIGMA), followed by 3 rounds of washes with PBS with 0.5% BSA. Cells were then stained with DAPI, flow sorted into 96 well plates containing 2.5ul of RLT plus lysis buffer (Qiagen), collecting only viable singlets, but not excluding cells based on size or granularity. Plates were then frozen and stored at −80C until processing.

### Library preparation

#### Single cells - scM&T-seq

Sorted single cells were processed as previously described. Briefly, mRNA capture and sequencing was carried out using G&T-seq (Macaulay et al., 2015). gDNA was purified using AMPure XP beads and then processed using scBS-seq (Clark et al., 2017).

scRNA-seq libraries were sequenced on Illumina Hiseq4000 instruments using 75bp paired-end reads and pooling 384 cells per lane. scBS-seq libraries were sequenced using 150bp paired-end reads pooling 48 cells per lane.

#### Bulk and sorted blood cell types - RRBS

RRBS libraries were prepared from isolated DNA as described before (Stubbs et al., 2017). The final amplified libraries were purified, assessed for quality and quantity using High-Sensitivity DNA chips on the Agilent Bioanalyzer and sequenced using 75 bp paired-end protocol on a HiSeq 2000 instrument (Illumina). For this project we extended the data generated for the Stubbs et al study by: 1) generating extra RRBS samples coming from older mice but the same tissues (Liver, Lung, Cortex and Heart), extending the age range of the Babrahm study to 105 weeks maximum, and 2) additional RRBS runs were generated on new tissues and cel types, ages ranging from 14 weeks to 105 weeks). This second batch also contains sorted blood cells leveraged in this study (B-cells, CD4+ T-cells, and CD8+ T-cells), additionally we generated data on cerebellum, hind muscle, kidney, testes as well as intestinal stem cells, and lung macrophages.

### External datasets

In addition to the newly generated data, the following publicly available datasets were downloaded from GEO, processed as described below and included in the analysis: GSE93957 (Stubbs et al., 2017); GSE80672 (Petkovich et al., 2017); GSE60012 (Reizel et al., 2015); GSE121141 (Meer et al., 2018); GSE120137 (Thompson et al., 2018); SRA344045 (Gravina et al., 2016).

### Data processing

#### Single cells - scM&T-seq

scM&T-seq data was processed as described previously (DNAme: (Angermueller et al., 2016); RNA: (Linker et al., 2019)). scBS-seq data was processed as described previously; briefly, reads had 6bp removed on their 5’-ends to remove the random primed portion of the reads, and were also adapter- and quality-trimmed using Trim Galore (v0.6.7; options --clip_r1 6). Trimmed reads were aligned to the bisulfite converted GRCm38 mouse genome using Bismark v0.22.3 in singleend mode (options: --non_directional) (Krueger and Andrews, 2011). Methylation calls were extracted after duplicate reads had been removed (deduplicate_bismark). scRNA-seq data was processed as described previously, briefly: the RNA-seq reads were adapter and quality-trimmed using Trim Galore (v0.6.7), and aligned to the GRCm38 mouse genome build using STAR (2.7.1a) (Veeneman et al., 2015), in two-pass mode. Expression quantification was performed leveraging FeatureCounts available in Subread 1.6 (Liao et al., 2019, 2014), and based on ENSEMBL version 96 (Cunningham et al., 2022).

#### Single cell quality control and normalisation

scM&T-seq data was quality controlled pr level. On the DNAme side We removed cells with a low read depth (removing cells with < 1M reads), removed cells with a low unique number of mapping reads (cells with < 50,000 uniquely aligned reads are removed), and high non-CpG methylation levels (cells with non-CpG meth >20% are removed). This left a total of 853 high quality cells. For downstream analysis we mapped all CpGs to the forward strand and removed non-CpG or ambiguous methylation calls.

Quality control of scRNA-seq data was performed using the SCATER package (McCarthy et al., 2017). Cells were retained for downstream analysis if they had at least 150,000 counts from endogenous genes, at least 1,500 genes with non-zero expression, less than 90% counts came from the top 100 highest expressed features and less than 15% mitochondrial reads. After quality control, 981 out of 1,055 blood single-cells were considered for downstream analysis. For further analyses, expression counts were SCRAN normalised into counts per million and log transformed (log(normCPM+1)).

During our analysis we found that a proportion of cells had very high expression values for haemoglobin genes (Hbb-bt, Hbb-bs, Hba-a1, and Hba-a2), leveraging the expression counts of these genes we estimated the number of red blood (RB) cells contaminating our expression level. The expression of the four genes formed distinctive peaks, around no expression, around 5 reads per gene per cell and around 10 reads per gene per cell. We combined the information over the marker genes and defined cells with on average 5 reads per marker per cell as having 1 RB cell as contamination (348 cells), and if on average more than 10 reads per marker per cell was found we defined it as 2 contaminating RB cells (69).

#### Cell type annotation

We leveraged a combined de novo and reference-based mapping setup to annotate cells to cell types. Specifically, for the de novo we leveraged shared-nearest neighbour (SNN) from Seurat, as input we used the first 10 PCs, derived from the top 2,000 highly varying genes that are expressed in at least 1% of the cells, the SNN clustering resolution was set to 0.5. This yielded 12 clusters. In parallel we performed a reference-based cell annotation, to do so we took the bulk RNA-seq data from the haemopedia resource (de Graaf et al., 2016). The resource contains Bulk RNA-seq data of 57 different flow sorted mouse blood cell types, originating from 13 different cell lineages, and are derived from healthy mice. The raw data was reprocessed with the same pipeline as our single cell data, see above. From this we simulated single cells leveraging Splatter (Zappia et al., 2017), expression counts were taken from the bulk data, other parameters needed for the simulation were derived from the actual sc-RNAseq data. By leveraging SingleR (Aran et al., 2019) we subsequently annotated the individual cells to the best matching simulated single cells from Heamopedia and annotated them to the relevant cell type. Lastly, we combine the information from SNN and SingleR and do a majority vote per cluster to assign final cell types per cluster and thereby cells. We leveraged the cell type annotation derived from RNA also in the DNAme analyses. Given the low cell numbers we focused the cell type specific analysis to B cells, CD4+ T cells (EffCD4T, MemCD4T, NveCD4T, RegT), and CD8+ T cells (MemCD8T, NveCd8T).

### Bulk RRBS data processing and normalisation

RRBS and WGBS datasets were processed as described previously (Stubbs et al., 2017; von Meyenn, 2022), aligning reads to the bisulfite converted GRCm38 mouse genome using Bismark v0.22.3 (Krueger and Andrews, 2011). After mapping we transformed the DNAme calls to the forward direction and removed non-CpG or ambiguous methylation calls.

We performed per tissue (liver and blood) and dataset (Petkovich; Reizel; Meer; Thompson; Gravina; Babraham_p1, Babraham_p2, Babraham_p3) quality control. To do so we selected sites that had at least 5X coverage in 80% of the samples, we dropped samples with high missingness rate (>25%) and subsetted per dataset tissue combination to sites present in all samples. Based on this set we did a PCA, again per dataset tissue combination, and dropped outliers on PC1 and PC2, we defined outliers as samples with scores higher (or lower) than mean plus (or minus) two times SD on each of the two PCs.

### UMAP on expression and DNAme

To get an overview of the global structures in both the gene expression data and the DNAme data we used UMAP (McInnes et al., 2018). On expression we used the top 1,000 most highly expressed genes and derived form that PCs, and took the top 15 to make the UMAP plots (Fig 1c & 1d). On DNAme we joined the information on DNAme levels on promoters and enhancers, given the sparsity we removed regions with observations in less than 80% of the cells. Next we used the “imputePCA()” function from the missMDA (Josse and Husson, 2016) package to impute the missing information. Subsequently we selected the 25% most variable regions and made a UMAP on the 15 first PCs (Suppl Fig 1a & 1b).

### Tissue cell composition analysis

To test for effects of age on cell type composition we used propeller (Phipson et al., 2022), implemented in the Specle R package. We transformed the proportions using the logit function, leveraging “getTranssformedProps()” and used the “propeller.annova()” function to test for the effect of age, and correct for multiple testing.

### Differential expression analysis

We performed differential gene expression analysis leveraging MAST in the three major cell types (B-cell, CD4+ and CD8+ T-cells). Age, or epigenetic age, was used as a continuous variable and we treated, number of expressed genes, mouse ID, and Haemoglobin contamination as covariates. For T-cell analysis we also took sub-celltype proportion into account. For each of the three test cell types we filtered to: 1) protein coding genes, 2) genes expressed in at least 25% of the cells, selected the genes with a high biological variation (by measuring the “modelGeneVarByPoisson” function implemented in SCRAN (Lun et al., 2016) (var>1)), 3) and lastly only tested genes that are expressed in at least 5 cells in at least 2 distinct age groups. Additionally we tested for a binary effect of ageing between the ages lower then 101 weeks versus the 101 week old cells.

The MAST model implements a hurdle model that combines information from two tests, a continuous model (for cells with non-zero expression levels), and a binary model (testing expressed vs not expressed). The effect sizes of the continuous and discrete part of the model are then combined to give one final output. We filtered these to not have significant results (nominal P<0.05 but in opposite direction between the tests), and performed Storey’s Qvalue to account for multiple testing. Significance was defined at 10% FDR.

### Ageing affects numbers of genes expressed in mouse

To test the relation between age (and epigenetic age) and the number of genes expressed we used a linear model, implemented in R. For the test in our ageing mouse blood data we treated age as a binary variable (old: 101 weeks, young < 101 weeks) and corrected for number of expressed genes, cell types, and Haemoglobin contamination, unless otherwise specified.

For replication in the Tabula Muris data we mimicked the binary analysis testing ages lower or equal than 77 weeks versus ages higher than or equal to 101 weeks. Here again we corrected for the number of sequenced reads, and if appropriate corrected for sex. For the replication in the human PBMC data we matched these age criteria by selecting individuals below or equal to 55 years as young (roughly matching the 77 weeks in mouse), and assigned individuals over 64 years of age as old (roughly matching the 101 weeks age in mouse). In oneK1K we corrected the number of genes expressed for the number of sequenced reads, material batch, and sequencing pool.

### Differential methylation analysis

We performed the single cell differential DNAme analysis leveraging a generalised linear mixed effect model (GLMM) implemented in lme4 (Bates et al., 2015) and R. We leveraged a binomial link function to capture the binary nature of single cell DNAme data, and leveraged the random effect to capture the effects driven by mouse In this setup we tested for differentially methylated regions (DMRs) in enhancers and promoters, in the three major cell classes (B, CD4+ T and CD8+ T cells). We tested regions with DNAme information in more than 25% of the cells per cell type. The enhancer and promoter information was derived from UCSC.

To associate DNAme levels at CpG-island and repeat regions to ageing we combined all CpG sites in the relevant region category, covered in at least one cell of each of the donor mice. To assess the effect of age on the overall methylation levels we used a spearman rank test. To replicate this finding in the bulk data we selected these same sites, per category, and tested for the same effect in the sorted blood data, again using a spearman rank test.

### Modelling epigenetic age

To build an epigenetic clock that is able to deal with sparsity better as compared to standard elastic net regression models we leverage a direct distance-based model similar to the Trapp et al model and similar to genetic prediction models. We built these models based on the data of the Quality controlled RRBS samples as described above, the selection for CpGs is done independent from the QC procedure described above.

#### Building the expected age - methylation matrix

The first step we took was to build up an expected methylation versus age matrix. Todo so we combined published bulk RRBS blood (or liver) datasets, specifically new Babraham RRBS samples, GSE93957 (Stubbs et al., 2017), GSE80672 (Petkovich et al., 2017), GSE60012 (Reizel et al., 2015), GSE121141 (Meer et al., 2018), and GSE120137 (Thompson et al., 2018) and reprocessed the raw data as described above. Next we select CpG sites that are covered in at least 2 studies, have at most 25% missingness per covered dataset, and 33% maximum overall missingness in the combined study. In the blood dataset we included the pseudo bulked single cell data to inform the site selection on non-RRBS data. This leaves 366,250 CpG sites in blood and 753,296 CpG sites in liver.

After this initial selection we calculate per datasets and per CpG site a spearman correlation with age, we filter out CpGs with inconsistent correlation signs between datasets and combine this information by taking the average. For blood we again included the pseudobulked single cell data to filter for opposite effect signs, but they are not contributing to the eventual site selection. Subsequently we rank this list from highest to lowest absolute age correlating sites and prune this list for correlated sites. We do so by comparing the absolute correlation to age of sites within 5,000 bases from each other and if the age association is similar (absolute delta <0.1) then we only keep the top age associated site. The sites that are pruned away for the main age association but are kept as back-up sites, that can be used when the main associated site is not available in a test sample. For the independent age associate sites we then built up the methylation versus age matrix. To do so we model DNAme per site given age, using a binomial model, and take the dataset of origin as a 1-hot encoded fixed effect covariate along. We fit this model three times, once with linear age, once with log age and once with square root of age, to account for nonlinearities that age has with DNAme values. We selected the best transformation per site based on the residual variation after fitting the GLM. Leveraging these models per CpG we can calculate the DNAme values per age, and correct these for observed dataset effect, here we chose to only model one week higher and lower as compared to observations in our combined training set.

#### Predicting ages for new samples or cells

To model ages of new samples or cells we select the overlapping sites between the clock sites and a new sample and compare the DNAme levels, similar to the procedure outlined in Trapp et al. With one minor difference, we directly calculate the absolute difference between the given DNAme value and the expected values, and sum the log difference. This generalises the procedure proposed by Trapp et al to also work on continuous DNAme profiles as observed in bulk data.

Lastly, we used cross validation to select the number of age associated DNAme to us when predicting epigenetic ages. We tested 50,100,250,500,750,1000,1250,1500,1750,2000 sites for training, and found that 750 sites was optimal for both blood and liver.

#### Significance of the age deviation

To determine if an age prediction was significantly different from the expected chronological age we matched the real data to simulations matching the input cell and age. In detail, we select the predicted methylation profile matching the age of the cell, and select the sites that are covered in the cell of interest. From this we then simulate 1,000 single cells, which when combined match the expected methylation profile derived from the expected matrix for the exact same sites covered in a cell of interest. Subsequently we predict the ages of these simulated cells and can place the age prediction of the cell of interest in the distribution of random predictions. Based on this we can calculate if the real cell is an outlier in terms of age prediction, defined as less than 5% of the permuted predictions are higher, or lower then the real prediction (empirical FDR 5%).

### Enrichment analyses

To test for gene set enrichment analyses we leveraged g:Profiler (Reimand et al., 2007), when assessing enrichments for DNAme we mapped promoters and enhancers to the closest gene and did the enrichment via g:Profiler. When leveraging g:Profiler we made sure the backgrounds are matched to the tested genes, instead of the default whole genome background.

To test for CpG enrichments in the epigenetic clocks we leveraged a fisher exact test. We selected all considered sites in the clocks (scEpiAge-blood, scEpiAge-liverclock, or the Stubbs clock) as a background and counted the number of times a site would be within any of these categories (CpG islands, CGI shores, CGI shelves, CGI inter, FANTOM5 enhancers, Repeats, Gene bodies, lncRNAs, CDS, Introns, Exons, Intergenic regions, 3’ UTRs, 5’ UTRs, TES, TSS, Promoters, CGI Promoters, Non-CGI Promoters), stratifying for a selected clock site or a background site. The MM10 annotation was derived from UCSC.

## Acknowledgements

We thank all the members of the involved groups for helpful discussions. In particular, we thank Jonathan Clark (Head of the Biological Chemistry Facility at the Babraham Institute) for help and materials, Rachael Walker and the Babraham Flow Cytometry Team for help with cell sorting, the Wellcome Trust Sanger Sequencing Facility for assistance with high-throughput sequencing. We thank Fabian Springer, Astrid KM Stubbusch and Fatemeh Habibolahi, for help with the computational analysis, testing new pipelines and helping developing and writing scripts. We would also like to thank Daniel Bolland, Geoff Butcher, Tamir Chandra, Anne Corcoran, Melanie Eckersley-Maslin, Lucy Field, Ulrika C. Frising, Colin Gilbert, Joana Guedes, Irene Hernando-Herráez, Jon Houseley, Fiona Kemp, Amy MacQueen, Klaus Okkenhaug, Martyn Rhoades, Milou J. C. Santbergen, Marisa Stebegg, and Marc Veldhoen for help and support.

## Authors Contributions

SC, WR, and FvM designed the study; SC, AKS, TS, and FvM collected samples and performed experiments; FK performed processing of data; MJB, SC, SL, JAdS, AMH, SR, OS, WR and FvM analysed and interpreted data; MJB developed the sc DNAme model; MJB, SC, and FvM wrote the draft manuscript; All authors commented, edited, and approved the final manuscript.

## Ethics approval

Animal experiments were performed according to the UK Animals (Scientific Procedures) Act 1986 and approved by the Babraham Research Campus Animal Welfare and Ethical Review Body.

## Accession numbers

Code: https://github.com/EpigenomeClock/AgingClock_v2

## Supplementary material

Attached

