## Supplementary material for "Single cell DNA methylation ageing in mouse blood"

*Bonder et al*

**Supplementary Tables** 2

**Supplementary Figures** 3

### Supplementary tables

**Suppl Table 1**: Details of collected peripheral blood samples from mice spanning ages from 10 to 101 weeks

**Suppl Table 2**: Number of genes expressed changes with age in Tabula Muris and OneK1K

**Suppl Table 3**: Ageing associated genes

**Suppl Table 4**: Ageing associated genes enrichments

**Suppl Table 5**: Age-related DNAme changes in both enhancers and promoters

**Suppl Table 6**: Details of all bulk data sets included

### Supplementary Figure 1

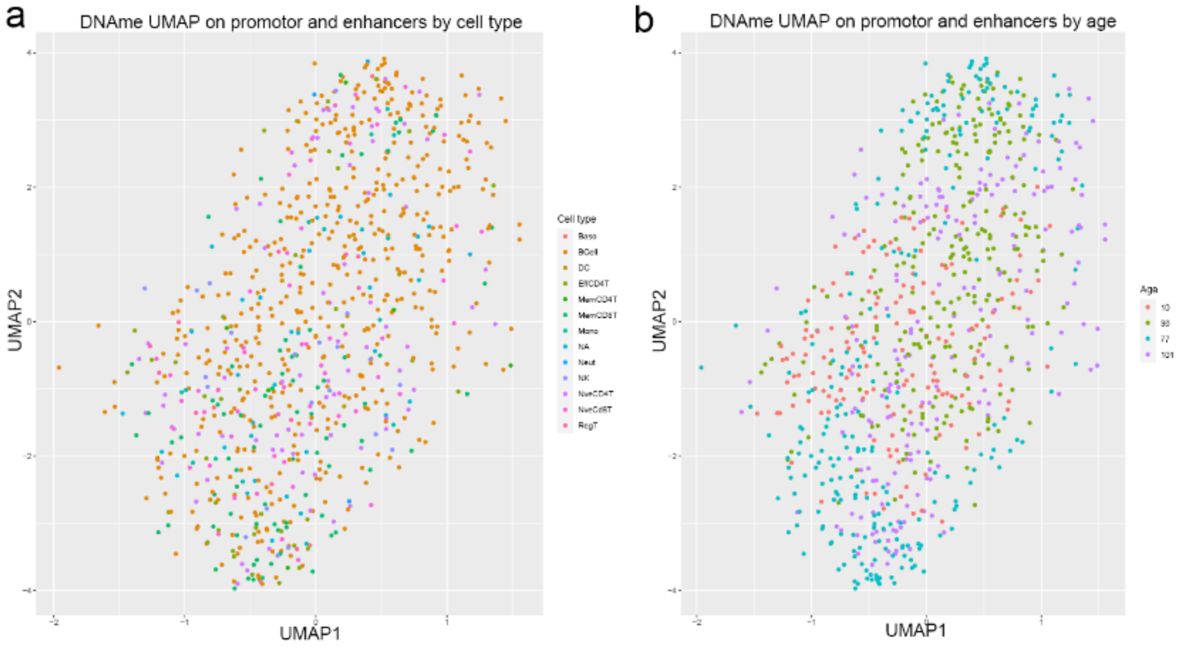


***Suppl Figure 1: DNAme did not show clear separations by age, cell type or animal.*** *Exploratory UMAP figures of the single cell DNAme data at enhancers and promoters:* ***a****) Cell type annotated UMAP, and* ***d****) UMAP annotated by chronological age.*

### Supplementary Figure 2

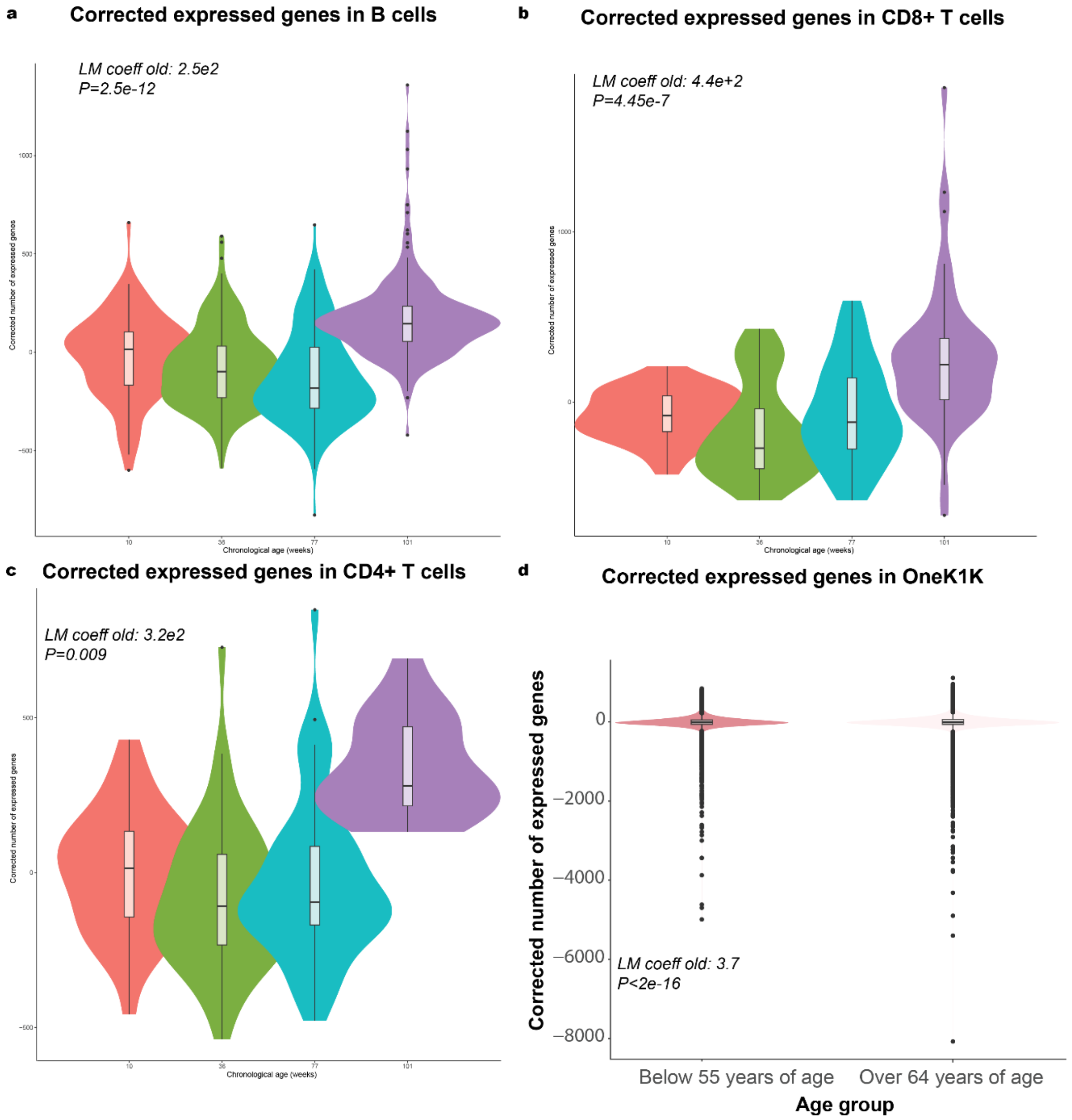


***Suppl Figure 2: Number of genes expressed per major cell type.*** *Violin plots showing the number of genes expressed/detected in each cell:* ***a-c)*** *Shown are the results for the major cell types analysed, namely* ***a)*** *B-cells,* ***b)*** *CD8+ T-cells, and* ***c)*** *CD4+ T-cells.* ***d)*** *Shown are the differences in a large human PBMC cohort, the OneK1K dataset (Yazar et al., 2022). To reflect the mouse analysis, we did split the humans data by age into samples below 55 years (roughly corresponds to below 77 weeks of age in mouse) and above 64 years (roughly corresponds to over 101 weeks of age in mice).*

### Supplementary Figure 3

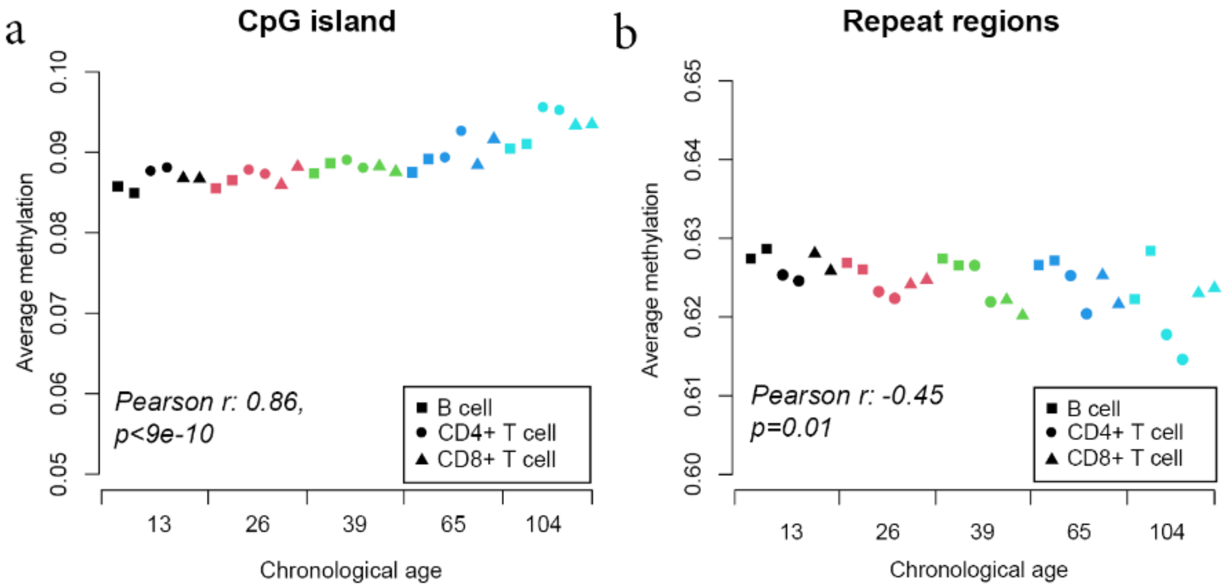


***Suppl Figure 3: scDNAme changes in the major cell types analysed.*** *Average single cell DNAme levels in* ***a)*** *CGIs and* ***b)*** *repeat regions. Cells are coloured by cell type (blue B-cell, orange CD4+ T-cell, green CD8+ T-cell) and ordered by age. Y-axis are scaled to the max and min values found in the presented data.*

### Supplementary Figure 4

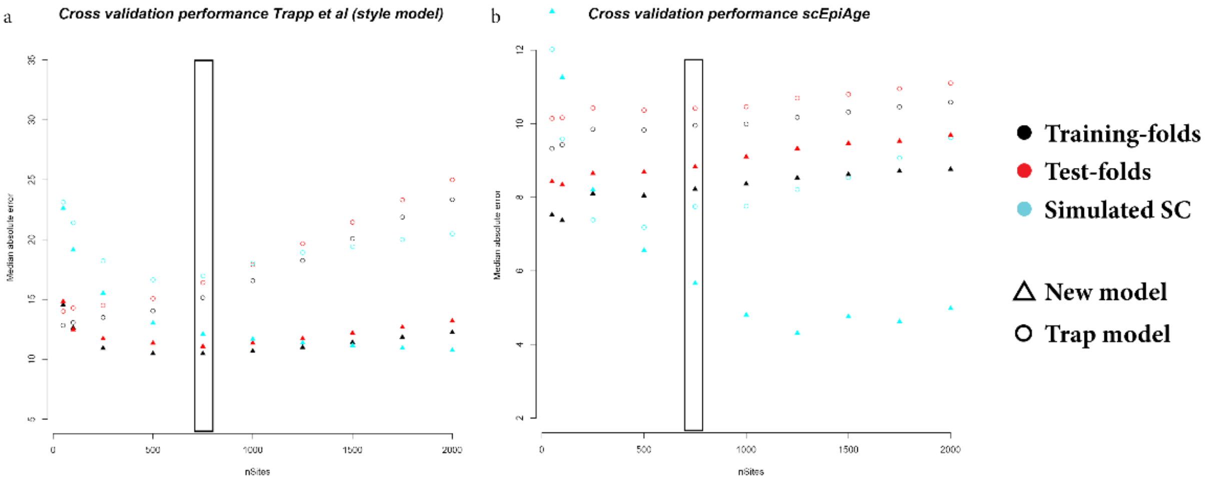


***Suppl Figure 4**: **Training and test performance during cross validation. a-b)** Error in training, testing as well as left out single cell data during cross validation optimising the number of sites included in the model. In **a)** the performance of model building using the Trapp et al setup, in **b)** the model setup in the scEpiAge setup.*

### Supplementary Figure 5

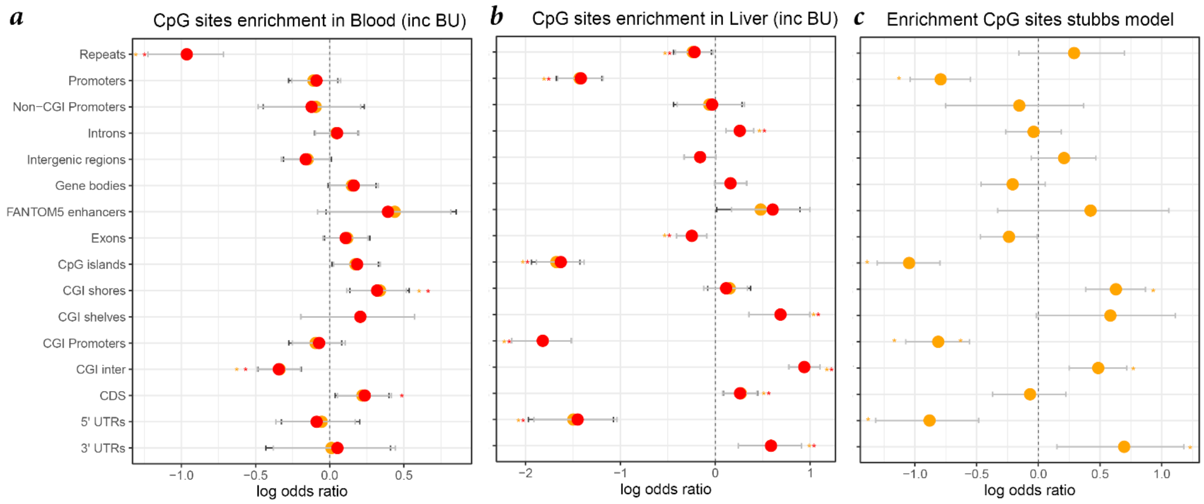


***Suppl Figure 5: Genomic enrichments of sites and back-up sites of the scEpiAge model.*** *We assessed the genomic enrichment of the sites (and back-up sites) selected in the* ***a)*** *scEpiAge blood model and* ***b)*** *scEpiAge liver model, and compared them to the sites selected for the* ***c)*** *Stubbs et. al clock (Stubbs et al., 2017).*

### Supplementary Figure 6

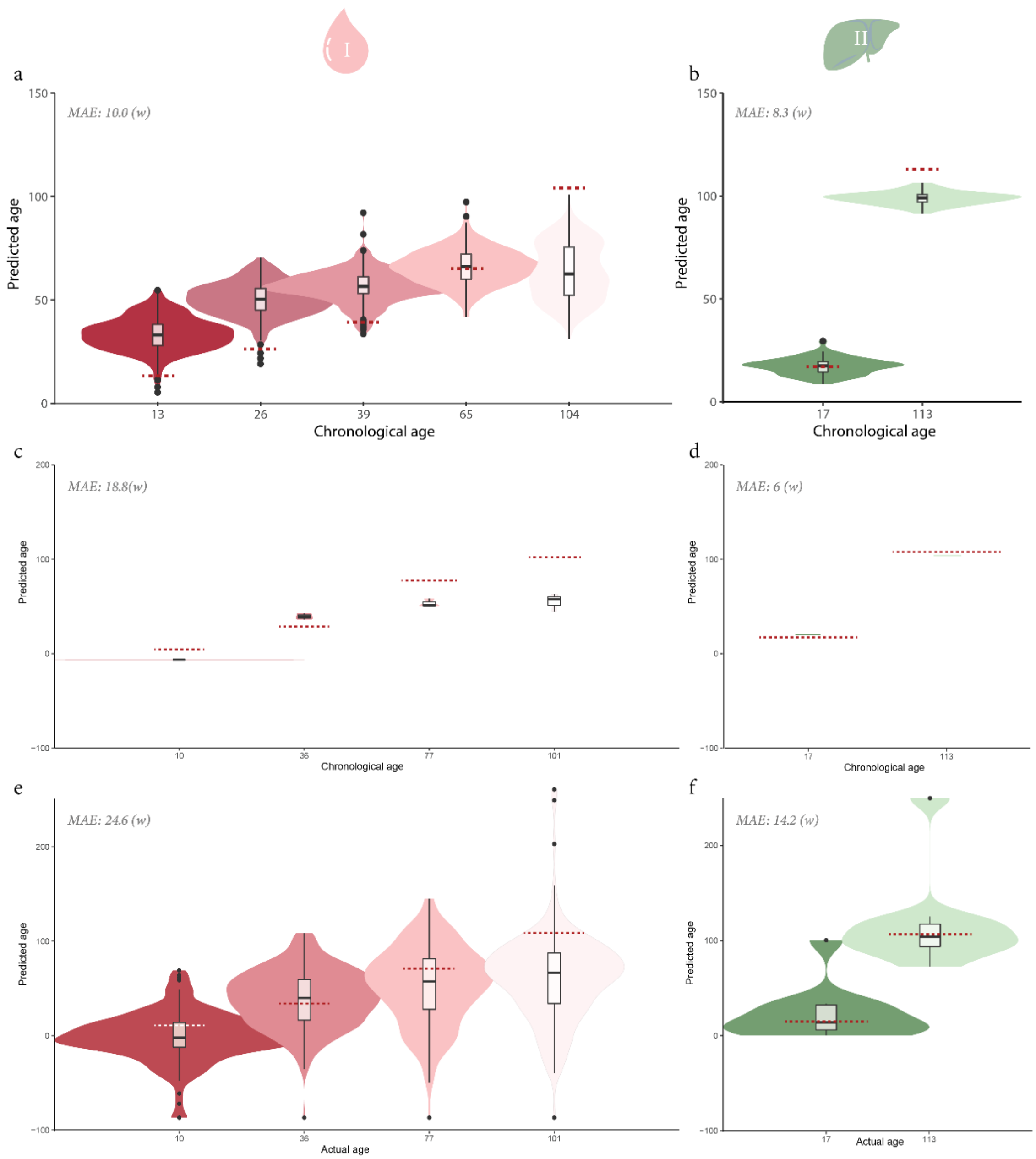


***Suppl Figure 6: Performance of the Trapp et. al model on scDNAme data.*** *We used the model described by Trapp et al. (Trapp et al., 2021) and assessed the performance of the model on simulated **a**) blood or **b**) liver single cells, pseudo bulked real **c)** blood and **d**) liver single cell data, and single cell **e**) blood and **f**) liver data.*

### Supplementary Figure 7

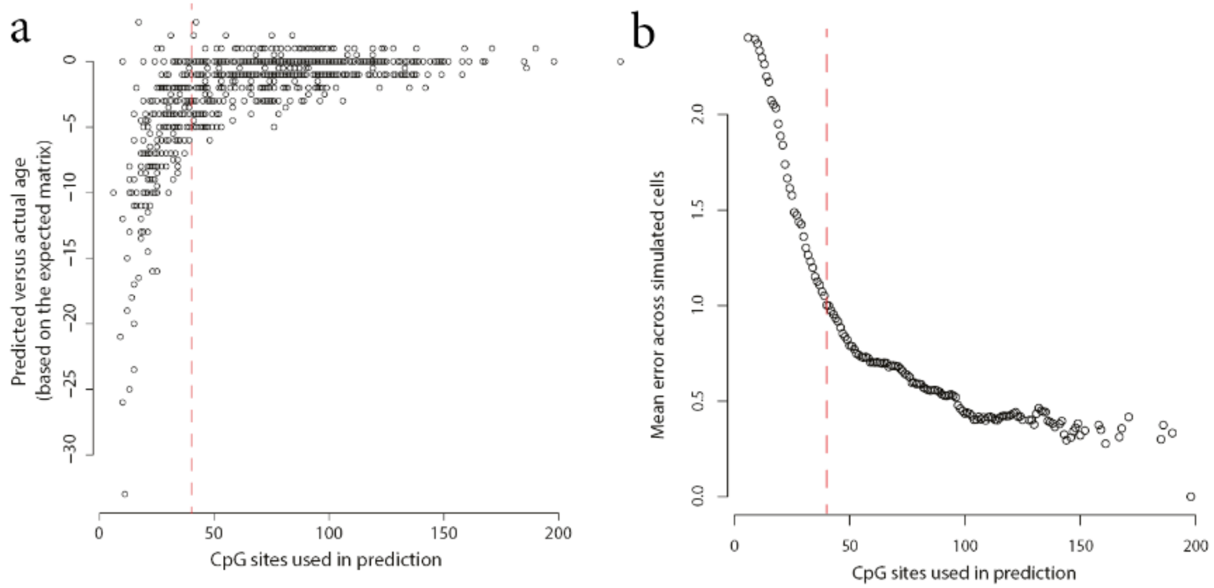


***Suppl Figure 7: Minimum overlapping sites needed for accurate scEpiAge predictions. a)*** *We calculated the difference between actual and predicted age in relation to the number of CpG sites used in prediction in simulated cells. For the expected data we matched the sites based on the actual cell.* ***b)*** *Additionally, we plotted the Average error of all simulated cells in relation to the number of CpG sites used in prediction.*

### Supplementary Figure 8

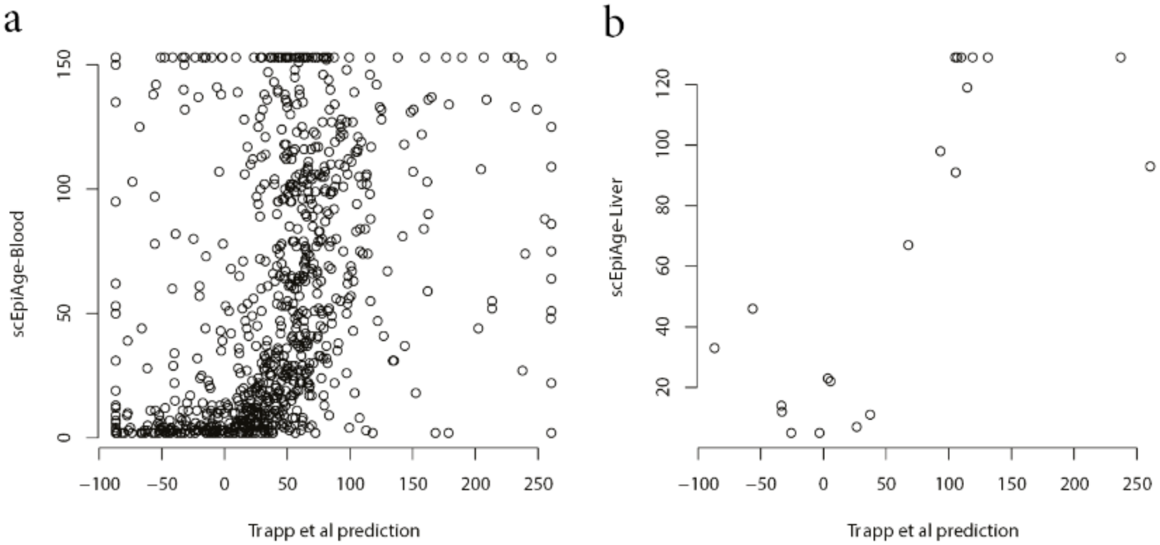


***Suppl Figure 8: Comparison between scEpiAge model and Trapp et al. model.*** *We used scEpiAge and the model from Trapp et al. (Trapp et al., 2021) to estimate the epigenetic age based on the DNAme data of all single cells. Shown are the correlation between both models for **a**) blood and **b**) liver single cell DNAme data.*
